# Modelling of synaptic interactions between two brainstem half-centre oscillators that coordinate breathing and swallowing

**DOI:** 10.1101/2021.05.04.442535

**Authors:** Pavel Tolmachev, Rishi R. Dhingra, Jonathan H. Manton, Mathias Dutschmann

## Abstract

Respiration and swallowing are vital orofacial motor behaviours that require the coordination of the activity of two brainstem central pattern generators (r-CPG, sw-CPG). Here, we use computational modelling to further elucidate the neural substrate for breathing-swallowing coordination. We progressively construct several computational models of the breathing-swallowing circuit, starting from two interacting half-centre oscillators for each CPG. The models are based exclusively on neuronal nodes with spike-frequency adaptation, having a parsimonious description of intrinsic properties. These basic models undergo a stepwise integration of synaptic connectivity between central sensory relay, sw- and r-CPG neuron populations to match experimental data obtained in a perfused brainstem preparation. In the model, stimulation of the superior laryngeal nerve (SLN, 10s) reliably triggers sequential swallowing with concomitant glottal closure and suppression of inspiratory activity, consistent with the motor pattern in experimental data. Short SLN stimulation (100ms) evokes single swallows and respiratory phase resetting yielding similar experimental and computational phase response curves. Subsequent phase space analysis of model dynamics provides further understanding of SLN-mediated respiratory phase resetting. Consistent with experiments, numerical circuit-busting simulations show that deletion of ponto-medullary synaptic interactions triggers apneusis and eliminates glottal closure during sequential swallowing. Additionally, systematic variations of the synaptic strengths of distinct network connections predict vulnerable network connections that can mediate clinically relevant breathing-swallowing disorders observed in the elderly and patients with neurodegenerative disease. Thus, the present model provides novel insights that can guide future experiments and the development of efficient treatments for prevalent breathing-swallowing disorders.

**Key points:**

- The coordination of breathing and swallowing depends on synaptic interactions between two functionally distinct central pattern generators (CPGs) in the dorsal and ventral brainstem.
- We model both CPGs as half-centre oscillators with spike-frequency adaptation to identify the minimal connectivity sufficient to mediate physiologic breathing-swallowing interactions.
- The resultant computational model(s) can generate sequential swallowing patterns including concomitant glottal closure during simulated 10s stimulation of the superior laryngeal nerve (SLN) consistent with experimental data.
- *In silico*, short (100 ms) SLN stimulation triggers a single swallow which modulates the respiratory cycle duration consistent with experimental recordings.
- By varying the synaptic connectivity strengths between the two CPGs and the sensory relay neurons, and by inhibiting specific nodes of the network, the model predicts vulnerable network connections that may mediate clinically relevant breathing-swallowing disorders.

## 1 INTRODUCTION

Swallowing and breathing are motor behaviours necessary for survival. Swallowing is essential for feeding and drinking, whereas breathing controls vital pulmonary gas exchange (oxygen uptake and excretion of carbon dioxide). Although these motor behaviours serve different biological functions, they both use a common oropharyngeal passage and thereby share a variety of upper airway muscles to control respiratory airflow (Bartlett Jr (1989), Dutschmann and Paton (2002b), Dutschmann and Paton (2002a), Dutschmann et al. (2014)) or the transportation of food or fluids from the oral cavity to the oesophagus and stomach (Miller (1986), Sejdic et al. (2018), Allen (2012), Jean (2001)). Thus, the motor activity underlying breathing and swallowing needs to be tightly coordinated (Hårdemark Cedborg et al. (2009), Bolser et al. (2013), Bautista and Dutschmann (2014), Huff et al. (2020)).

Respiration and swallowing are motor activities which are generated and coordinated by anatomically overlapping central pattern generators (r-CPG, sw-CPG) in the brainstem (Bianchi and Gestreau (2009)). The r-CPG is anatomically organised into bilateral ponto-medullary columns of respiratory neurons (Alheid et al. (2004)). However, more recent studies revived the notion that generation of the eupneic three-phase respiratory motor pattern consisting of inspiration, postinspiration and expiration also requires synaptic interactions with the dorsal respiratory group (Berger (1977), Hilaire et al. (1990), Dhingra et al. (2019a), Dhingra et al. (2019b), Dhingra et al. (2020)). The sw-CPG is located in the caudal medulla oblongata and is subdivided into the dorsal swallowing group located in the nucleus of the solitary tract (NTS) (Car et al. (1979), Kalia and Mesulam (1980), Kessler and Jean (1985)) and the ventral swallowing group, dorsomedial to the nucleus ambiguus (Kessler and Jean (1985); Ezure et al. (1993); Sugiyama et al. (2011), Fuse et al. (2019)). More recent data suggest that the ventral swallowing group may functionally overlap with the postinspiratory complex (Toor et al. (2019), Pitts et al. (2021)). While breathing is unconditional, and must be maintained throughout a lifetime, swallowing is a conditional motor act which is initiated by adequate sensory inputs that are mediated by the superior laryngeal nerves (Miller and Loizzi (1974), Jafari et al. (2003)) or behavioural commands (Mosier and Bereznaya (2001)).

The laryngeal adductor muscles that constrict the vocal folds are integral for both swallowing and respiration (Bolser et al. (2006), Bianchi and Gestreau (2009), Davenport et al. (2011), Pitts et al. (2013)). Partial laryngeal adduction controls the duration and strength of expiratory airflow (Harding (1984), Bartlett Jr (2011), Dutschmann and Paton (2002b), Dutschmann and Paton (2002a), Bautista and Dutschmann (2014)) and protects the lower airways (trachea, lungs) from aspiration of foreign material, in particular, during sequential (rhythmic) swallowing (Medda et al. (2003)). A previous study identified that the pontine Kölliker-Fuse nucleus mediates protective laryngeal adduction during swallowing (Bautista and Dutschmann (2014)).

The precise synaptic mechanism underlying the generation of sequential or single swallows, as well as the synaptic connectivity between neuron populations of the sw-CPG and r-CPG underlying coordination of breathing and swallowing are not completely understood. A main obstacle is the difficulty to identify functional neuroanatomy of the synaptic interactions that coordinate breathing and swallowing activity within the intact network since r-CPG and sw-CPG overlap anatomically and functionally (Bianchi and Gestreau (2009)). Computational modelling studies can be effective tools to study the connectivity and function of neural networks (Molkov et al. (2017), Lindsey et al. (2012)). During the past two decades, Hodgkin-Huxley-based models of synaptic mechanisms underlying respiratory rhythm generation and the formation of a three-phase respiratory motor pattern were the main interest in the field (Rybak et al. (1997), Rybak et al. (2004), Smith et al. (2007), Rubin et al. (2009), Molkov et al. (2013), Molkov et al. (2014), Molkov et al. (2017)). While the putative network response to sensory stimuli, such as the Hering-Breuer reflex of the pulmonary stretch receptors (Molkov et al. (2013)) or hypoxia/hypercapnia (Molkov et al. (2014)) were modelled previously, we are only aware of a single publication that modelled the interaction of two different motor CPGs during respiratory entrainment (Potts et al. (2005)). Moreover, only a conceptual model for breathing-swallowing interaction has been published previously (Bolser et al. (2013)).

In the present study, we extend previous computational models of the respiratory CPG to include the interactions with the swallowing CPG, which is modelled as a conditionally active half-centre oscillator (HCO). The intrinsic neuronal dynamics are simplified, and the equations underlying the Matsuoka neural oscillator are utilised (Matsuoka (2011)). Thus, the present model focuses on the connectivity between the neural populations rather than their biophysical properties. The same limited number of model parameters is used for the modelling of r-CPG function, however, the connectivity between respiratory neuronal populations is largely adapted from previous r-CPG models (Smith et al. (2007), Rybak et al. (2004), Rubin et al. (2009)) to generate a three-phase motor pattern of breathing (Richter (1982), Smith et al. (1991), Rubin et al. (2009)). The present computational model is supported by and compared to the experimental data obtained in a perfused brainstem preparation. Accordingly, the present model can generate sequential swallowing with concomitant glottal closure during tonic SLN stimulation (10s). Moreover, the present model can reproduce the swallowing phase response curve as well as the generation of single swallows in response to short (100-250ms) stimulation of the SLN. Finally, an extensive sensitivity analysis of the modelled network connectivity predicts vulnerable synaptic pathways that can be linked to the clinically relevant swallowing disorder such as aspiration (Yagi et al. (2017)), seen in patients with various neurodegenerative diseases (Takizawa et al. (2016), Kalia (2003), Chouinard (2000), Prosiegel et al. (2004), Heemskerk and Roos (2011)) and the elderly (Farpour et al. (2018), Sura et al. (2012), Finiels et al. (2001)).

## 2 MATERIALS AND METHODS

### 2.1 Experimental data

Experimental protocols were approved by and conducted with strict adherence to the guidelines established by the Animal Ethics Committee of the Florey Department of Neuroscience and Mental Health, University of Melbourne, Melbourne, Australia.

Experiments were performed in juvenile (17-21 days postnatal) Sprague-Dawley rats of either sex using the in situ arterially perfused brainstem preparation as described previously (Paton (1996), Dutschmann et al. (2000)). Briefly, rats were anesthetised by inhalation of isoflurane (2-5%) until they reached a surgical plane of anesthesia. Next, rats were transected sub-diaphragmatically and immediately transferred to an ice-cold bath of artificial cerebrospinal fluid (aCSF; in mM: 125 [*NaCl*_2_], 3 [*KCl*], 1.25 [*KH*_2_*PO*_4_], 1.25 [*MgSO*_4_], 24 [*H*_2_*CO*_3_], 2.5 [*CaCl*_2_]) for decerebration. Next, the heart and lungs were removed. The phrenic nerve was isolated for recording, and the descending aorta was isolated for cannulation. Next, the cerebellum was removed. Finally, the vagus and superior laryngeal nerves were isolated for recording and stimulation, respectively.

The preparation was then transferred to a recording chamber. The aorta was quickly cannulated with a double-lumen catheter. The preparation was then reperfused with carbogenated (95%/5% *pO*_2_/*pCO*_2_), heated (31 C°) aCSF using a peristaltic pump (Watson-Marlow). The phrenic and vagal nerves were mounted on suction electrodes to record the fictive respiratory and swallowing motor patterns. Motor nerve potentials were amplified (400×), filtered (1-7500 Hz), digitised (30 kHz) via a 16-channel differential headstage (Intan RHD2216), and stored on an acquisition computer using an Open-Ephys acquisition system (Rev. 2, Siegle et al. (2017)). Within minutes, apneustic respiratory contractions resumed. A single bolus of NaCN (0.1 mL, 0.1% (w/v) NaCN) provided an excitatory chemosensory drive to convert the apneustic pattern to a eupneic three-phase respiratory motor pattern. Finally, the perfusion flow rate was adjusted to fine-tune the preparation to generate a stable stationary rhythm.

To evoke single or sequential swallows, we stimulated the superior laryngeal nerve (SLN). The threshold to evoke swallowing was determined by applying 10s trains (20 Hz, 0.1 ms pulse width) at increasing current amplitudes (10-80 *µ*A). Next, we measured the sequential swallowing evoked by SLN stimulation at twice the threshold current. To measure the SLN phase resetting curve, single swallows (n=100 swallows) were evoked at random intervals between 20 – 25 s (20 Hz, 250 ms burst duration, 0.1ms pulse width, 2 × threshold current) such that SLN stimulation was evoked at every phase of the respiratory cycle.

### 2.2 Data analysis of experimental data

The processing of the recordings of the vagus and phrenic nerves was implemented using python programming language, utilising functions implemented in scipy package (Virtanen and Contributors (2020)). First, the mean value of the recordings was subtracted. Afterwards, the recordings were filtered with a cascade of two 2^*nd*^ order band-pass Butterworth filters (signal.butter, cut-off frequencies: 300 Hz, 3 kHz). For this and the following filterings, a filter was applied once forwards, once backwards, to avoid introducing phase delays (signal.filtfilt function). The SLN stimulus artifacts were removed by linear interpolation of the signals around the stimulus time, negating the immediate short-term effects caused by the stimulus. The absolute value was taken, and the signals were further filtered using a 2^*nd*^ order low-pass Butterworth filter (12 Hz cut-off). The resultant vagus nerve activity resembled a respiratory airflow, and both signals were more amenable for further analysis. In the end, the signals were decimated by a factor of 100 by cascading the function signal.decimate twice, each time applying an anti-aliasing filter and reducing the signal lengths by a factor of 10. The final sample rate of the signals was 300 Hz.

In the present study, we evaluate a phase response curve (PRC) of the breathing-swallowing circuit -– the dependence of phase shift of the respiratory oscillations on the phase at which the short SLN stimulus was applied. To obtain the PRC, the continuous recording was separated into shorter sequences that contain a single SLN stimulation (Fig. 1). For each sequence, the recorded phrenic nerve activity (PNA) was additionally filtered using a 2^*nd*^ order low-pass filter (`signal.butter`, cut-off freq. = 1 Hz) to retain only the primary respiratory harmonics (the typical respiratory frequencies are 0.2-1 Hz). The Hilbert transform was applied afterwards to have an initial estimate of the signals’ phase (protophase, Kralemann et al. (2008)). The mean of the resulting analytic signal was subtracted, and the protophase was obtained. Using a linear fit (`optimize.minimize`) of the extracted protophase before the stimulus, the frequency of oscillations *ω*_0_ was estimated. At the next step, a refined estimation of phase *ϕ* was obtained, using the procedure described in (Kralemann et al. (2008)). A linear regression on the phase was performed, using the fit *ϕ*_*b*_ = *ω*_0_ *t* + *b*_1_ before and *ϕ*_*a*_ = *ω*_0_*t* + *b*_2_ after the SLN stimulus (with *ω*_0_ fixed), and the free coefficients *b*_1_ and *b*_2_ were extracted. The phase shift was obtained according to Δ*ϕ* = (*b*_1_ - *b*_2_) (See Fig. 1).

**FIGURE 1.**
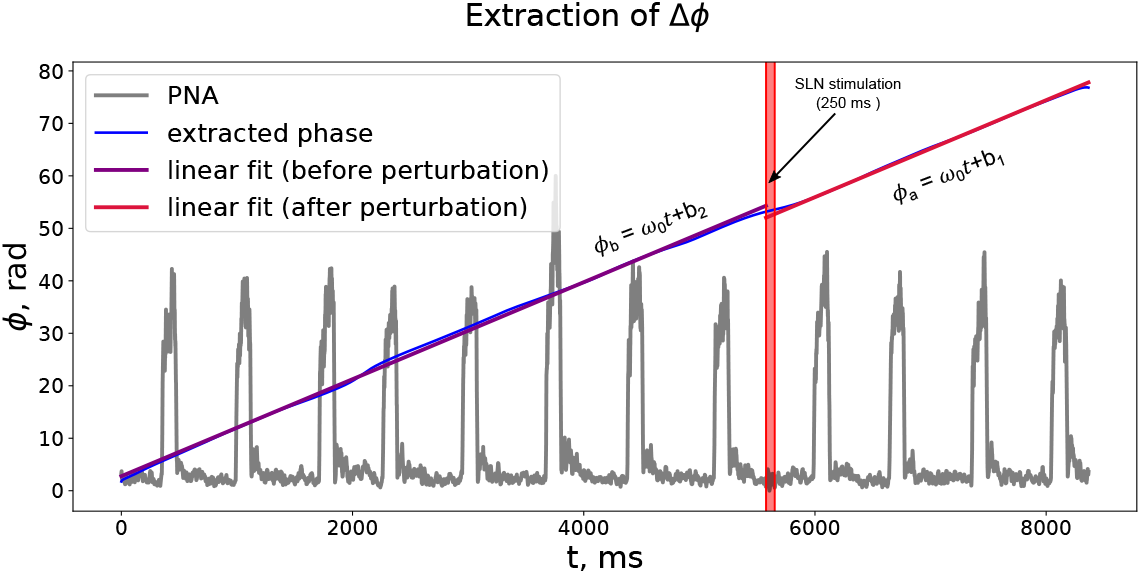
A short stimulation of the superior laryngeal nerve (SLN) affects the respiratory rhythm. The phase change after the short (250 ms) SLN stimulation (depicted by red) is reflected in the processed experimental recording of the phrenic nerve activity (PNA). The phase of the respiratory rhythm was estimated before and after the stimulation, and the resulting phase shift was extracted. See text for the details.

### 2.3 Model parameters

Previous studies that modelled r-CPG function incorporated Hodgkin-Huxley style biophysical properties of respiratory neurons (Smith et al. (2007), Rubin et al. (2009), Molkov et al. (2013), Molkov et al. (2014), Molkov et al. (2017)). However, such models are prone to overfitting (Del Negro et al. (2018)). Instead of using Hodgkin-Huxley style description, we model neuronal populations with simpler equations underlying the Matsuoka neural oscillator (Matsuoka (2011)); that allows for the model of r-CPG and sw-CPG to function with a reduced set of parameters:

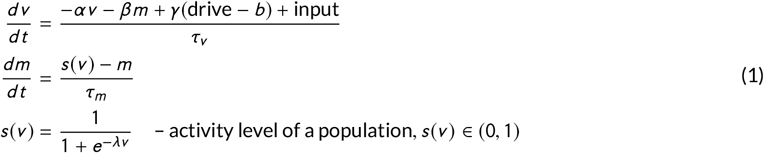

In these equations, the dimensionless variable *v* is analogous to the average membrane potential of a whole neural population. The variable *m* mediates the firing rate (or spike frequency) adaptation and acts on a much slower timescale than the variable *v*. To account for the correct timescale of the dynamics and the excitability of the neuronal populations, we use *α* = 1, *β* = *γ* = 100, *b* = 0.2 *τ*_*v*_ = 0.5 ms, *λ* = 0.3. The parameter *τ*_*m*_, which controls the timescale of spike-frequency adaptation, has distinct values for the different neuronal populations (discussed later), and the details are summarised in 3). The parameters are chosen to match the timescale of respiratory phase durations (inspiration, postinspiration, late expiration) of experimental recordings.

The term ‘input’ in the equations incorporates synaptic input from the other nodes of the neuronal populations of the r-CPG or sw-CPG, as well as the sensory input from the SLN (applicable only to the Sensory Relay neurons, see later). The input to a *j* ^*t*^ *h* neural population is modelled by the following equation:

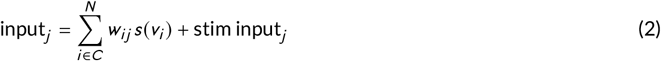

Where *C* is the set of neural nodes projecting to the *j* ^*th*^ neuron. The connectivity parameters *w*_*ij*_ between the nodes are manually adjusted to reproduce realistic swallowing and breathing interactions. The gradual development of the model (see below, 2) allowed for adjusting connectivity parameters step by step, and the connectivity of a simpler model served as a reference point for the connectivity of a subsequent model. The baseline level of activity of a neuronal population is controlled by the parameter ‘drive’ which physiologically corresponds to the tonic excitatory drive from the other brain regions (discussed later).

### 2.4 Successive development of model connectivity

Using the equations for the neuronal populations defined above, we developed the network connectivity in 4 successive steps (Fig. 2). The initial model (Model 1; Fig. 2) established the basic connectivity: a half-centre oscillator in the r-CPG generates biphasic respiratory activity (inspiration (‘Insp’) and expiration (‘Exp’)), and a conditionally active half-centre oscillator for the sw-CPG (‘Sw_1_’, ‘Sw_2_’ populations) produces a sequence of rhythmic swallows in response to a simulated 10s sensory input from the SLN. The premotor ‘Sw_1_’ neurons contribute to the vagus nerve discharge. The ‘Sensory Relay’ neurons, whose activity indicates the presence of the sensory stimulus, are comprised of excitatory and inhibitory neuronal populations. The excitatory subpopulation of the ‘Sensory Relay’ neurons provide drive for the sw-CPG, while the inhibitory one suppresses the activity of the inspiratory neurons in the r-CPG during ongoing swallowing. In the initial model, a 10s sensory input from the SLN (Fig. 3) evokes the swallowing related bursts of activity in the vagus nerve output for the larynx that resembles the pharyngeal phase of swallowing (for details see Jean (2001), Bautista et al. (2014b), Lang (2009)). The large value of spike-frequency adaptation time constant of the ‘Sensory Relay’ neurons is responsible for a slowly declining frequency of the sequential swallows. The simulated SLN stimulation effectively suppresses inspiratory activity during the 10s period of swallowing.

**FIGURE 2.**
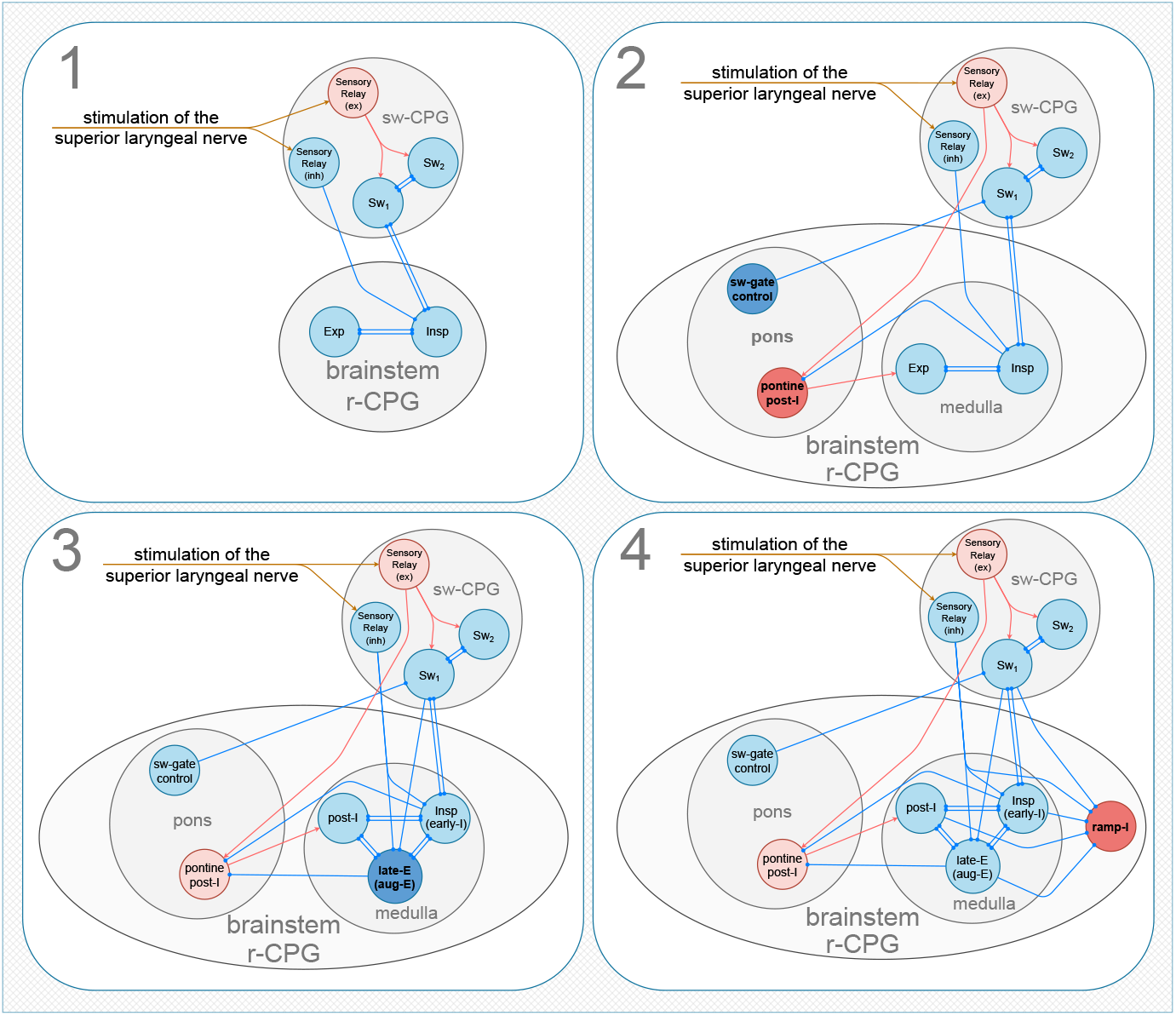
Progression of models of the interaction between the swallowing and the respiration circuits. The red and blue nodes denote excitatory and inhibitory neural populations, respectively. Blue connections capped with circles depict one-sided inhibitory projections, while the red arrows denote the excitatory ones. Starting from the two half-centre oscillators for respiratory and swallowing CPGs (r-CPG, sw-CPG) (1), we add pontine populations; one tonically inhibits sw-CPG, another one controls the glottal closure reflex and fires in phase with respiratory ‘post-I’ neurons (2). Further, we extend the r-CPG to account for the three-phase rhythm by introducing ‘late-E’ neural population (3). Lastly, to account for the correct ramping shape of the inspiration activity in the phrenic nerve, we introduce ‘ramp-I’ population (4).

**FIGURE 3.**
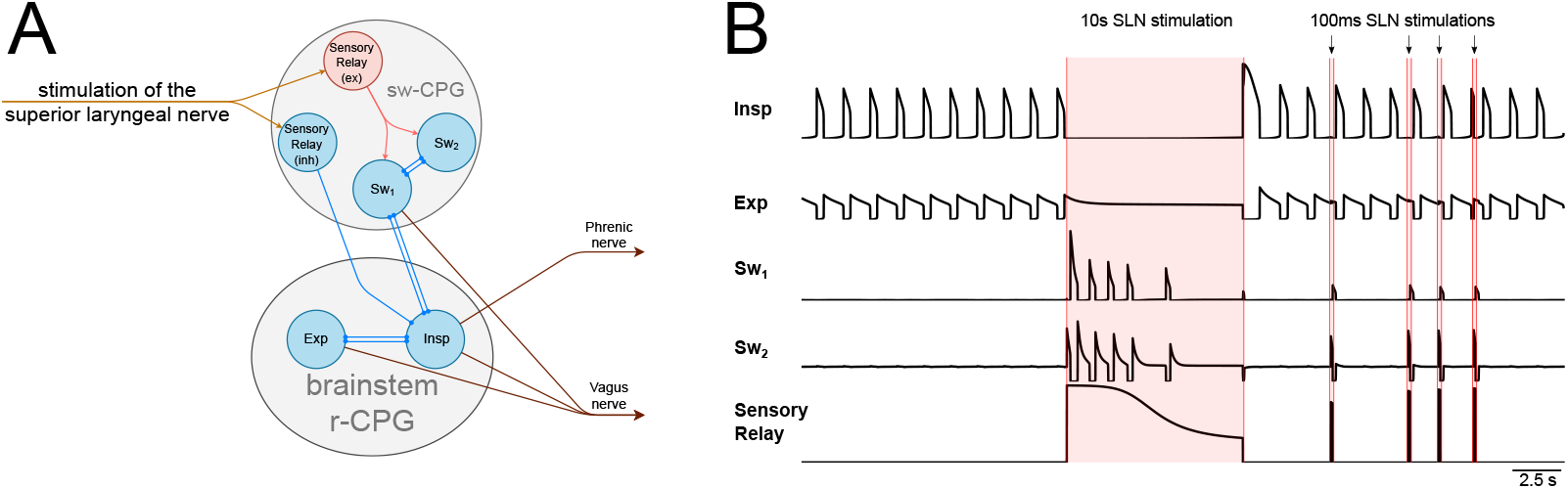
(**A**) The core model underlying the breathing-swallowing interaction. The respiratory CPG, to the first approximation, is modelled as a half-centre oscillator, producing the sustained inspiration-expiration alternation. The swallowing CPG is modelled as a conditionally active half-centre oscillator. (**B**) The swallowing half-centre oscillator produces sequential bursts of activity during the simulated 10 s sensory stimulus (long red shading), while the respiratory activity is silenced. The short sensory stimulus evokes a single swallow in the swallowing CPG, which interferes with the ongoing respiratory activity.

The next step of model evolution (Model 2; Fig. 2) implemented the separation of the r-CPG into the medullary and pontine compartments, to accommodate experimental findings that the pons contains neuronal populations that control the gating of the sw-CPG (Bautista and Dutschmann (2014)). In addition, an excitatory postinspiratory neuronal population was introduced into the pons to account for the tonic postinspiratory activity that mediates laryngeal constriction during sequential swallowing (Bautista and Dutschmann (2014)).

The next stage of the model evolution (Model 3, Fig. 2) incorporates an additional expiratory neuronal population to the r-CPG (‘late-E’) to generate a three-phase respiratory motor pattern, as the hallmark of previous r-CPG model simulations (Smith et al. (1991), Rybak et al. (1997), Rybak et al. (2004), Rubin et al. (2009), Molkov et al. (2013), Molkov et al. (2014), Molkov et al. (2017)). The core of the r-CPG is thus modelled as a ring of three mutually inhibiting populations of inspiratory (‘Insp’), postinspiratory (‘post-I’) and late-expiratory population (‘late-E’, active during the second half of the expiration, *E*_2_).

In the final model (Model 4; Fig. 2) we incorporate an additional premotor ‘ramp-I’ population of neurons to account for a eupneic respiratory motor activity that requires a ramping discharge pattern of the simulated phrenic nerve activity (PNA) and vagus nerve activity (VNA) (John and Paton (2003), Harris et al. (2017)). No intrinsic neural mechanisms have been discovered that account for the firing rate acceleration followed by the abrupt cessation of activity, thereby, this shape of discharge was hypothesised to originate from the inhibitory (primarily local) connections (Harris et al. (2017)). In the final model, the activity levels of the neural populations to simulate the outputs of PNA and VNA are modelled as a linear combination of activities of ‘ramp-I’, ‘pontine post-I’ and ‘Sw_1_’ with the coefficients 0.9, 0.75 and 0.6 correspondingly. The connectivity parameters between the populations are presented in Table 1.

**TABLE 1.**
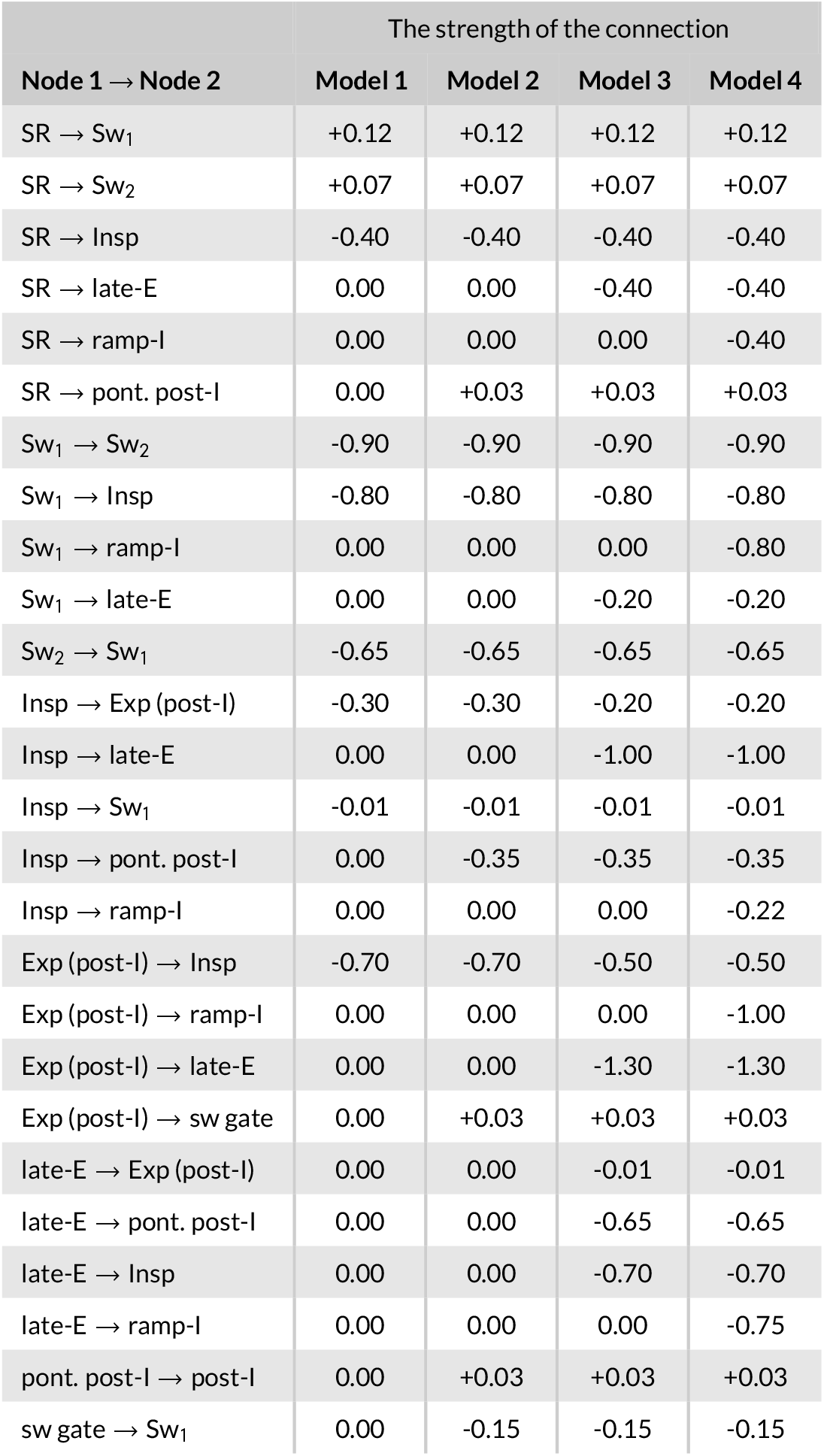
Connectivity description

| Node 1 → Node 2 | The strength of the connection |  |  |  |
| --- | --- | --- | --- | --- |
|  | Model 1 | Model 2 | Model 3 | Model 4 |
| SR → Sw<sub>1</sub> | +0.12 | +0.12 | +0.12 | +0.12 |
| SR → Sw<sub>2</sub> | +0.07 | +0.07 | +0.07 | +0.07 |
| SR → Insp | -0.40 | -0.40 | -0.40 | -0.40 |
| SR → late-E | 0.00 | 0.00 | -0.40 | -0.40 |
| SR → ramp-I | 0.00 | 0.00 | 0.00 | -0.40 |
| SR → pont. post-I | 0.00 | +0.03 | +0.03 | +0.03 |
| Sw<sub>1</sub> → Sw<sub>2</sub> | -0.90 | -0.90 | -0.90 | -0.90 |
| Sw<sub>1</sub> → Insp | -0.80 | -0.80 | -0.80 | -0.80 |
| Sw<sub>1</sub> → ramp-I | 0.00 | 0.00 | 0.00 | -0.80 |
| Sw<sub>1</sub> → late-E | 0.00 | 0.00 | -0.20 | -0.20 |
| Sw<sub>2</sub> → Sw<sub>1</sub> | -0.65 | -0.65 | -0.65 | -0.65 |
| Insp → Exp (post-I) | -0.30 | -0.30 | -0.20 | -0.20 |
| Insp → late-E | 0.00 | 0.00 | -1.00 | -1.00 |
| Insp → Sw<sub>1</sub> | -0.01 | -0.01 | -0.01 | -0.01 |
| Insp → pont. post-I | 0.00 | -0.35 | -0.35 | -0.35 |
| Insp → ramp-I | 0.00 | 0.00 | 0.00 | -0.22 |
| Exp (post-I) → Insp | -0.70 | -0.70 | -0.50 | -0.50 |
| Exp (post-I) → ramp-I | 0.00 | 0.00 | 0.00 | -1.00 |
| Exp (post-I) → late-E | 0.00 | 0.00 | -1.30 | -1.30 |
| Exp (post-I) → sw gate | 0.00 | +0.03 | +0.03 | +0.03 |
| late-E → Exp (post-I) | 0.00 | 0.00 | -0.01 | -0.01 |
| late-E → pont. post-I | 0.00 | 0.00 | -0.65 | -0.65 |
| late-E → Insp | 0.00 | 0.00 | -0.70 | -0.70 |
| late-E → ramp-I | 0.00 | 0.00 | 0.00 | -0.75 |
| pont. post-I → post-I | 0.00 | +0.03 | +0.03 | +0.03 |
| sw gate → Sw<sub>1</sub> | 0.00 | -0.15 | -0.15 | -0.15 |

We implemented multiple sources of tonic drives for the r-CPG and sw-CPG neuronal populations in the final model: the ‘Arousal drive’, the ‘Descending pontine drive’ for the medullary respiratory circuit, and the ‘Chemo-drive’ (see Fig. 4). In accordance with experimental findings (Sato et al. (2011), Sato et al. (2016)), the ‘Arousal drive’ controls the excitability of the sw-CPG. The ‘Descending pontine drive’ predominantly regulates excitability of ‘post-I’ neurons in the medullary compartment of the r-CPG (Dutschmann and Herbert (2006), Rybak et al. (2004)). Lastly, the ‘Chemo-drive’ projects to ‘late-E’ populations but also provides excitation to the medullary inspiratory neurons. The specific settings for the tonic drives are summarised in Table 2.

**FIGURE 4.**
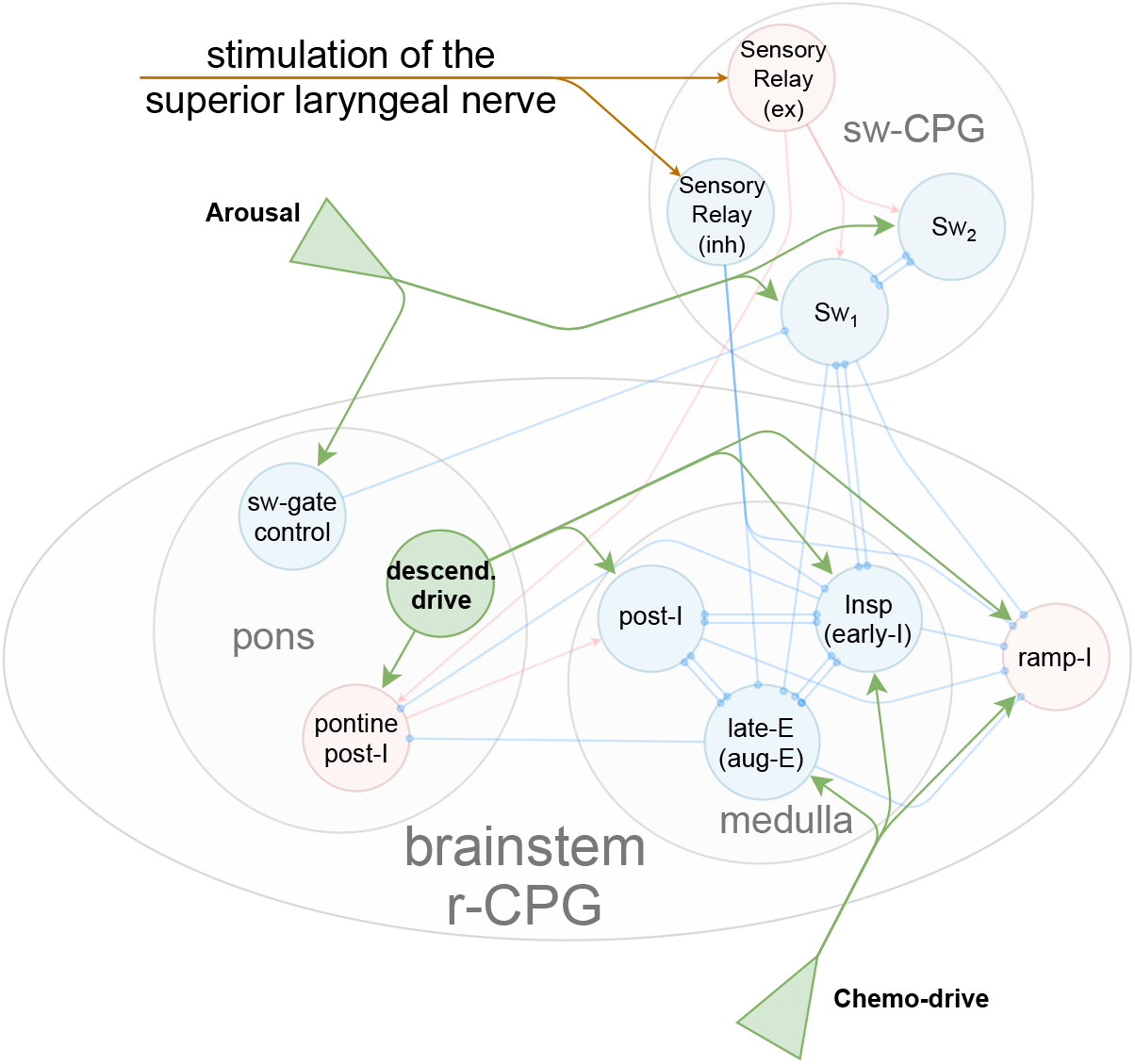
The full model of the respiratory and the swallowing CPGs depicted with the tonic drives. The sources of the tonic drives are depicted by green shapes. The green arrows represent the input pathways. In our model we implement three sources of drives: 1) ‘Arousal drive’, controlling the excitability of swallowing neurons, 2) ‘Descending drive’ residing in the pons and controlling the excitability of the core of the r-CPG, 3) ‘Chemo-drive’, which adjusts its activity according to levels of O_2_, CO_2_ and pH. In the model, the effects of these drives are modelled as constant inputs in the right-hand side of the equations governing the dynamics of the neural nodes.

**TABLE 2.** Description of the excitatory drives to the populations

| Drive | Value |
| --- | --- |
| Chemo drive → Insp | 0.30 |
| Chemo drive → late-E | 0.45 |
| Chemo drive → pont. post-I | 0.35 |
| Wake → Sw<sub>1</sub> | **0.19\*** |
| Wake → Sw<sub>2</sub> | 0.35 |
| Wake → sw gate | 0.25 |
| Pons → Exp (post-I) | 0.35 |
| Pons → Insp | 0.05 |
| Pons → ramp-I | 0.05 |

\* the Wake drive to Sw<sub>1</sub> in the Model 1 is 0.16.

The timescale of spike-frequency adaptation of the modelled neuronal populations is controlled by the variable *τ*_*m*_. In the model, the spike-frequency adaptation of the neurons in sw-CPG is set to occur faster than in the neuronal populations of the medullary r-CPG. Such difference in the intrinsic properties is introduced to accommodate for the generation of a physiologically plausible number of swallows during simulated 10s SLN stimulation. Moreover, the ‘Sensory Relay’ and the ‘pontine post-I’ populations have much lower rates of spike-frequency adaptation than the medullary r-CPG populations, to account for the slower timescale of a decrementing laryngeal adductor activity during 10s SLN stimulation. The intrinsic properties of the neural groups are summarised in Table 3.

**TABLE 3.** Intrinsic parameters

| Population | $\tau_m$ |
| --- | --- |
| Insp | 5000 |
| ramp-I | 5000 |
| Exp (post-I) | 5000 |
| late-E | 5000 |
| Sw<sub>1</sub> | 1500 |
| Sw<sub>2</sub> | 1500 |
| Sensory Relay | 17500 |
| pont. post-I | 10000 |
| sw gate control | 5000 |

for all of the populations: $\alpha = 1, \beta = \gamma = 100, b = 0.2,$ $\tau_v = 0.5, \lambda = 0.3$

### 2.5 Modelling of a single swallow in response to short SLN stimulation in the sw-CPG

In physiological experiments using the perfused brainstem preparation, a short (100-250 ms) stimulation of the SLN evoked a single swallow after the termination of the stimulus, which caused modulation of the respiratory cycle duration (See Section 3 for details). To account for these experimental findings, we implemented the ‘Sensory Relay’ neurons exciting both ‘Sw_1_’ and ‘Sw_2_’ neuronal populations, which together comprise a swallowing half-centre oscillator. During eupneic breathing, the ‘Sw_2_’ neurons receive more arousal drive than the ‘Sw_1_’ neurons, leading to the inhibition of the ‘Sw_1_’ premotor population. In our model, a 100 ms SLN stimulation provides additional excitation to both ‘Sw_1_’ and ‘Sw_2_’ populations; however, the ‘Sw_2_’ neurons outcompete the ‘Sw_1_’ population, and, at the onset of the stimulus, the ‘Sw_2_’ neurons further inhibit the ‘Sw_1_’ population. After cessation of the 100ms SLN stimulation, the ‘Sw_1_’ premotor neurons escape the afferent inhibition and, with short latency, produce a single swallowing motor burst. This mechanism is further analysed and described using a phase space analysis (See Sec. 3).

## 3 RESULTS

The present computational model of breathing-swallowing interaction is based on equations for the Matsuoka neural oscillator. Initially, we model each CPG as a half-centre oscillator: the respiratory half-centre oscillator underlies the inspiratory-expiratory phase oscillations, while the swallowing half-centre oscillator generates a stimulus-dependent sequential pharyngeal swallowing activity (Fig. 3). Later, we progressively introduce additional neuronal populations (2) to account for the experimental data, both previously obtained and presented in this paper.

The final model (Fig. 4) can reproduce the eupneic three-phase motor pattern of respiration characterised by augmenting phrenic nerve activity and decrementing postinspiratory activity in the vagal nerve. During simulation of 10s synaptic input from the SLN forces the two reciprocally connected swallowing populations to compete and oscillate between active (‘Sw_1_’) and silent phases (‘Sw_2_’) resembling the activity during the pharyngeal phase of swallowing Jean (2001), Bautista et al. (2014b), Lang (2009)). The excitatory premotor subpopulation of ‘Sw_1_’ neurons drives the vagal motor output to the larynx. In parallel, ‘Sensory Relay’ neurons inhibit the inspiratory (‘Insp’) and late expiratory (‘late-E’) neuronal populations and arrest the three-phase respiratory cycle in the postinspiratory phase (‘post-I’). In addition, excitatory ‘Sensory Relay’ neurons project to the ‘pontine post-I’ neural population, which excites the medullary ‘post-I’ neurons and drive the vagal motor output to the larynx. The latter mediates the protective laryngeal adduction during sequential swallowing in accordance with the previously published data (Bautista and Dutschmann (2014)). Comparison of the model performance to experimental data shows that 10s electrical stimulation of the SLN elicits 5 to 14 swallows, while simulation of 10s SLN input triggers 5 to 7 swallows (see, Fig. 5).

**FIGURE 5.**
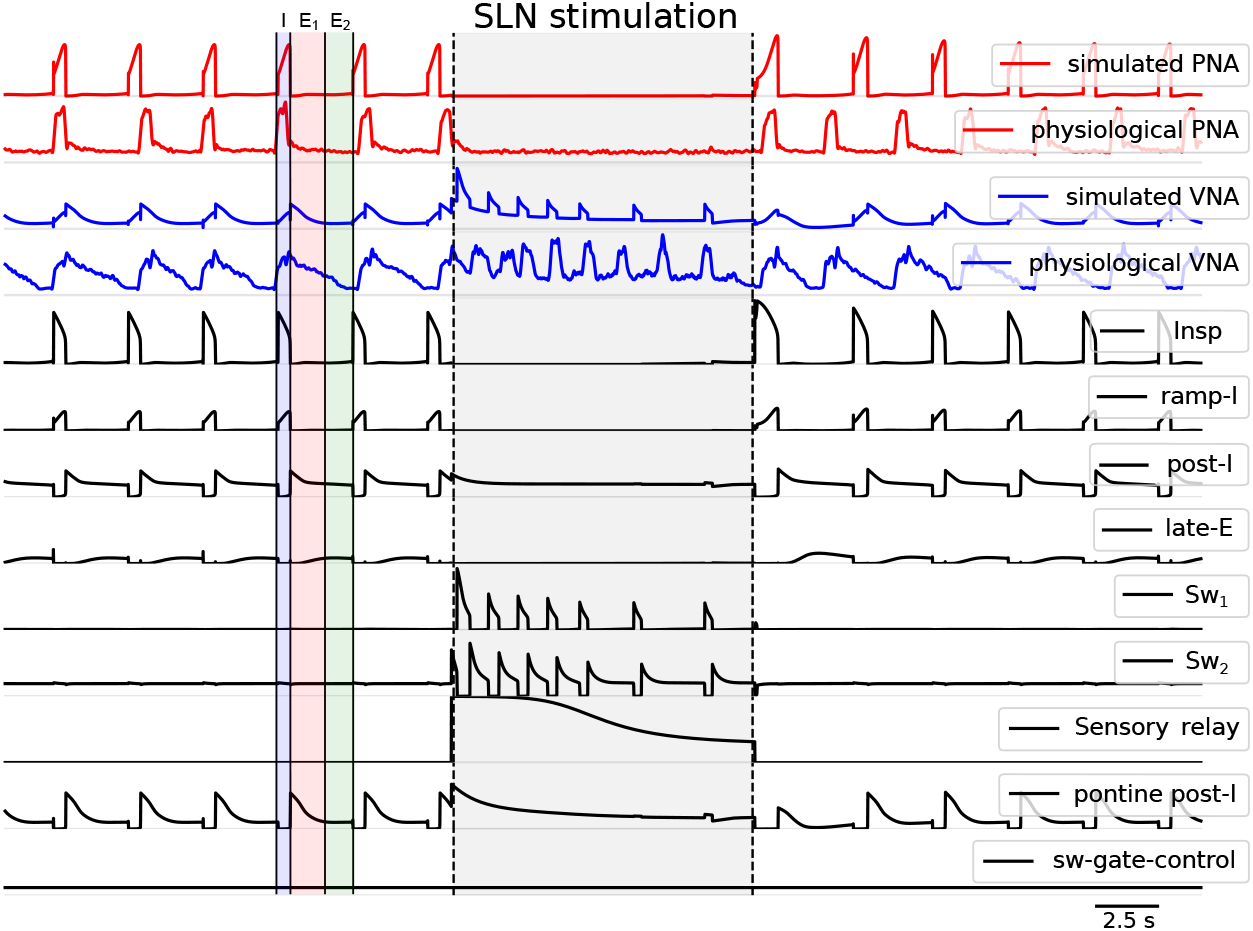
Comparison of the model output with the original recordings. The model reproduces the normal breathing with distinguished three phases. The comparison of experimental and simulated recordings from the activities of the phrenic and vagus nerve activities (PNA, VNA) is presented on the four upper traces. The rest of the traces show the activities of the individual neuronal populations. The model also qualitatively reproduces the effects of the superior laryngeal nerve (SLN) stimulation. The oscillations in r-CPG are silenced by the ‘Sensory Relay’ neurons, and the respiratory rhythm is locked in the postinspiratory phase. Meanwhile, the sensory neurons activate the sw-CPG, and the network produces swallowing-related oscillations, with the progressively decreasing frequency of swallows. During the sequential swallowing, the phasically active pontine ‘post-I’ population of neurons in the dorsolateral pons is responsible for the laryngeal adduction (glottal closure reflex). In the model, we assume that the phrenic nerve activity is driven solely by the ‘ramp-I’ neurons, whereas the vagus nerve activity is simulated as a combination of ‘pontine post-I’, ‘ramp-I’ and ‘Sw_1_’ populations with coefficients 0.75, 0.9 and 0.6 correspondingly.

The model also reproduces the experimental data of randomly applied short SLN stimulations in the perfused brainstem preparation. Similar to the experimental data, the simulated 100ms stimulus evokes a single swallowing-related burst, which causes timing-dependant modulation of the respiratory cycle length (Lewis et al. (1990), Oku and Dick (1992), for details see bellow and section 3.3; Fig. 1, Fig. 8).

### 3.1 Phase space analysis of the sw-CPG: transient dynamics of the swallowing half-centre oscillator in response to a short train of SLN stimulation

A short stimulation of the superior laryngeal nerve triggers a swallow which either delays or advances the phase of respiratory rhythm, depending on the timing of the SLN stimulation (Lewis et al. (1990), Oku and Dick (1992)). The mechanisms of the generation of a short-latency single swallow in response to short SLN stimulation could depend on the ion channel composition (HCN, BK and SK channels, Molineux et al. (2008)) of swallowing neuronal populations. Since our modelling approach aims to minimise parameterisation of biophysical properties of network neurons, we propose the generation of the short-latency swallowing response based on the network mechanism instead.

We implemented an asymmetric tonic excitatory drive (‘Arousal drive’) for the ‘Sw_1_’ and ‘Sw_2_’ neural populations and the inhibition of the ‘Sw_1_’ neuronal population by the descending tonic inputs from the pontine swallowing gate population (‘sw-gate control’). Taken together, these inputs provide a baseline excitation drive 0.16 for ‘Sw_1_’ and 0.35 for ‘Sw_2_’ neurons. Hence, during sensory inputs from the SLN, these settings render the ‘Sw_1_’ premotor neurons to be less excitable than the ‘Sw_2_’ population. In the model, the ‘Sw_1_’ neurons mediate the swallowing-related vagal motor output. To account for the physiologically observed short latency of swallow triggered by a short SLN stimulation and visible in the VNA, we implemented the connectivity of the sensory input to the swallowing populations so that the swallowing response starts from the ‘Sw_2_’ neurons. The required setting for the excitatory inputs from the ‘Sensory Relay’ neuronal population could then be determined by a phase space analysis. Figure 6 illustrates two different scenarios that discriminate the occurrence of a short-latency single swallow in response to a short SLN stimulation: in the first scenario, a short-latency swallowing response is observed after the stimulation, resembling the swallowing response in the experimental conditions; whereas in the second scenario, the swallowing response is immediate and concurrent with the stimulus, being unphysiological.

**FIGURE 6.**
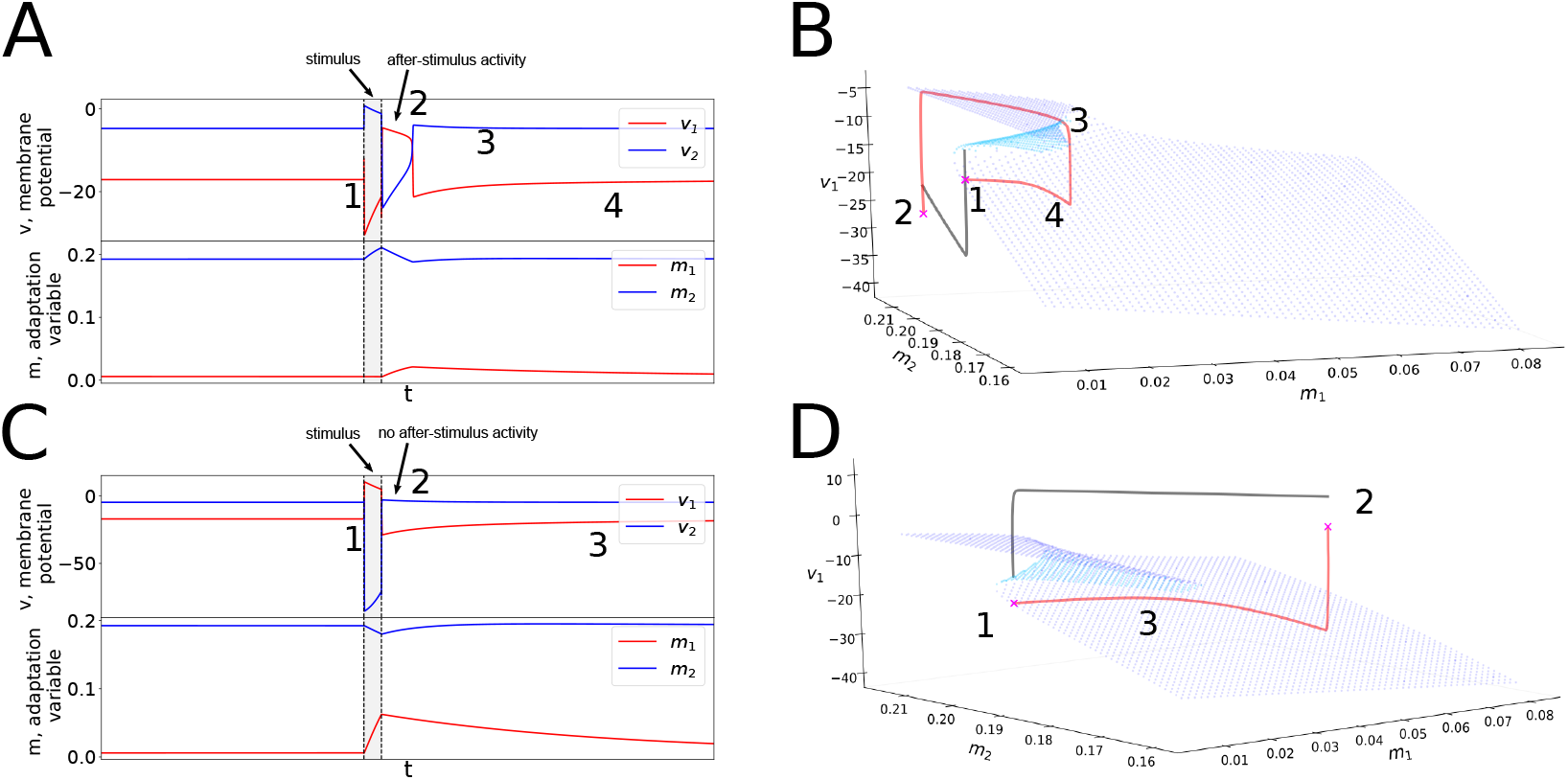
A short 100 ms stimulation of the superior laryngeal nerve (depicted by grey shadings) elicits a response in the sw-CPG. Here we isolate the swallowing CPG and provided an external stimulus from the ‘Sensory Relay’ neurons to study the mechanisms of the generation of the single swallow in the swallowing half-centre oscillator. Sub-figures (**A**) and (**C**) depict the traces of the half-centre oscillator: the membrane potential variables *v*_1_, *v*_2_ and adaptation variables *m*_1_ and *m*_2_; here, index ‘1’ denotes the premotor swallowing neurons. The swallowing premotor neurons are strongly inhibited in the absence of the external input, however, are readily excitable. The phase space analysis of the ongoing dynamics is depicted in the subfigures (**B**) and (**D**). Since the *v* -variables (fast subsystem) evolve on a much faster time scale than the *m* variables (slow subsystem), for each pair of the *m*-variables the equilibria points for the *v* -variables of the dynamics were calculated. Taken altogether, these fixed points form a cusp-like equilibrium surface of the fast subsystem. The slow subsystem governs the evolution of the whole system once the system reaches the stable submanifolds of the equilibrium surface. The stable submanifold of the surface is denoted by the navy blue colour, versus sky blue for the unstable equilibria. In both scenarios (**B**) and (**D**), the equilibria surface is the same. The difference between the two scenarios lays in the values of the stimulus supply to each population of the half-centre oscillator: in the first case, the stimulation strengths are (0.1, 0.05) to ‘Sw_1_’ (premotor) and ‘Sw_2_’ populations correspondingly, whereas in the second scenario the stimulation strengths are (0.15, 0.05), thereby in the latter case the ‘Sw_1_’ neuron inhibits ‘Sw_2_’ during the stimulus, however, returns to quiescence after the transient burst of activity.

Both scenarios differed only in the strengths of the sensory input to the swallowing populations. In the first scenario (Fig. 6 A, short-latency single swallow), the strengths of the sensory input to the ‘Sw_1_’ and ‘Sw_2_’ neurons required a setting of 0.10 and 0.05 correspondingly. In the scenario 2, a small increase of the input strength to the ‘Sw_1_’ population up to 0.15 prevents the occurrence of the normal swallowing response to the short SLN stimulation (See Fig. 6 C, the swallowing response is concurrent with the SLN stimulation).

To study the difference between these scenarios we performed a phase space analysis of the dynamics of the sw-CPG. The evolution timescales of *v* -variables and m-variables differ by three orders of magnitude, thereby the whole system can be subdivided into fast and slow subsystems. During the evolution, the *v*_1_ and *v*_2_ variables are almost functionally dependent, thereby one of the fast variables could be eliminated from the analysis, and the dynamics are effectively described in the three-dimensional phase space. For each fixed pair of slow *m*-variables (*m*_1_, *m*_2_) we calculated fixed points of the fast subsystem dynamics 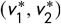. The resulting equilibrium surface described in (*m*_1_, *m*_2_, *v*_1_)-space has a cusp, which is crucial for the network dynamics (Fig. 6 B). In the global equilibrium (depicted by point 1 in the Fig. 6 A, B), the system stays on the lower part of the equilibrium surface, albeit close to the cusp. The location of the global equilibrium corresponds to the quiescent ‘Sw_1_’ neurons, since the value *s* (*v*_1_) is close to zero, although the system is readily excitable.

In the first scenario, the stimulus shifts the state of the system far from the lower sheet of the surface to the area with the overhanging upper sheet, and the state trajectory of the system jumps “vertically”, quickly reaching the upper part of the equilibrium surface. From here, it slowly slides down along the equilibrium surface until it reaches the other boundary of the cusp, and the single swallow terminates (Fig. 6 A, B, points 1-2-3).

Contrary to the first scenario, in the second scenario, the applied stimulus causes the system to jump to a state far from the upper sheet of the equilibrium surface. When the stimulus ceases, the system immediately relaxes back to the lower sheet of the surface (where ‘Sw_1_’ is quiescent), causing no transient after-stimulus activity (Fig. 6 C, D, points 1-2-3). In the model, the swallowing response is thereby determined by the cusp-like shape of the equilibrium surface of the fast subsystem, alongside the values of the sensory drives applied to the half-centre oscillator.

### 3.2 Model validation: deletion of the ponto-medullary synaptic interactions

Previously published data have shown that pharmacological inhibition of the pontine compartment of the r-CPG triggered apneusis (prolonged inspiratory phase and absence of postinspiratory motor discharge in VNA) and abolished the protective laryngeal adduction (Bautista and Dutschmann (2014)).This study also reported that pharmacological inhibition of the pons also initiated spontaneous swallows in the absence of the sensory input. The results of simulated inhibition of the pons are depicted in Fig. 7 B and the motor output is compared to the simulated eupneic activity (Fig. 7 A). The simulation of pontine inhibition is characterised by an increase in inspiratory phase duration (*T*_*i*_ from 0.41 ± 0.05 to 1.29 ± 0.05 s) and expiratory phase duration (*T*_*e*_ from 2.11 ± 0.05 to 3.55 ± 0.05 s)and the absence of postinspiratory discharge in the vagus nerve activity, reducing the rhythm to two phases. This phenomenon is explained by the absence of the descending pontine drive to medullary ‘post-I’ population, and thereby ‘late-E’ neurons suppress the activity of ‘post-I’ neurons. In addition, the present model features a pontine-dependent ‘sw-gate control’ population, and the removal of this population causes spontaneous swallows at the end of the inspiratory phase even in absence of the sensory input, in agreement with previous experimental data (Bautista and Dutschmann (2014)).

**FIGURE 7.**
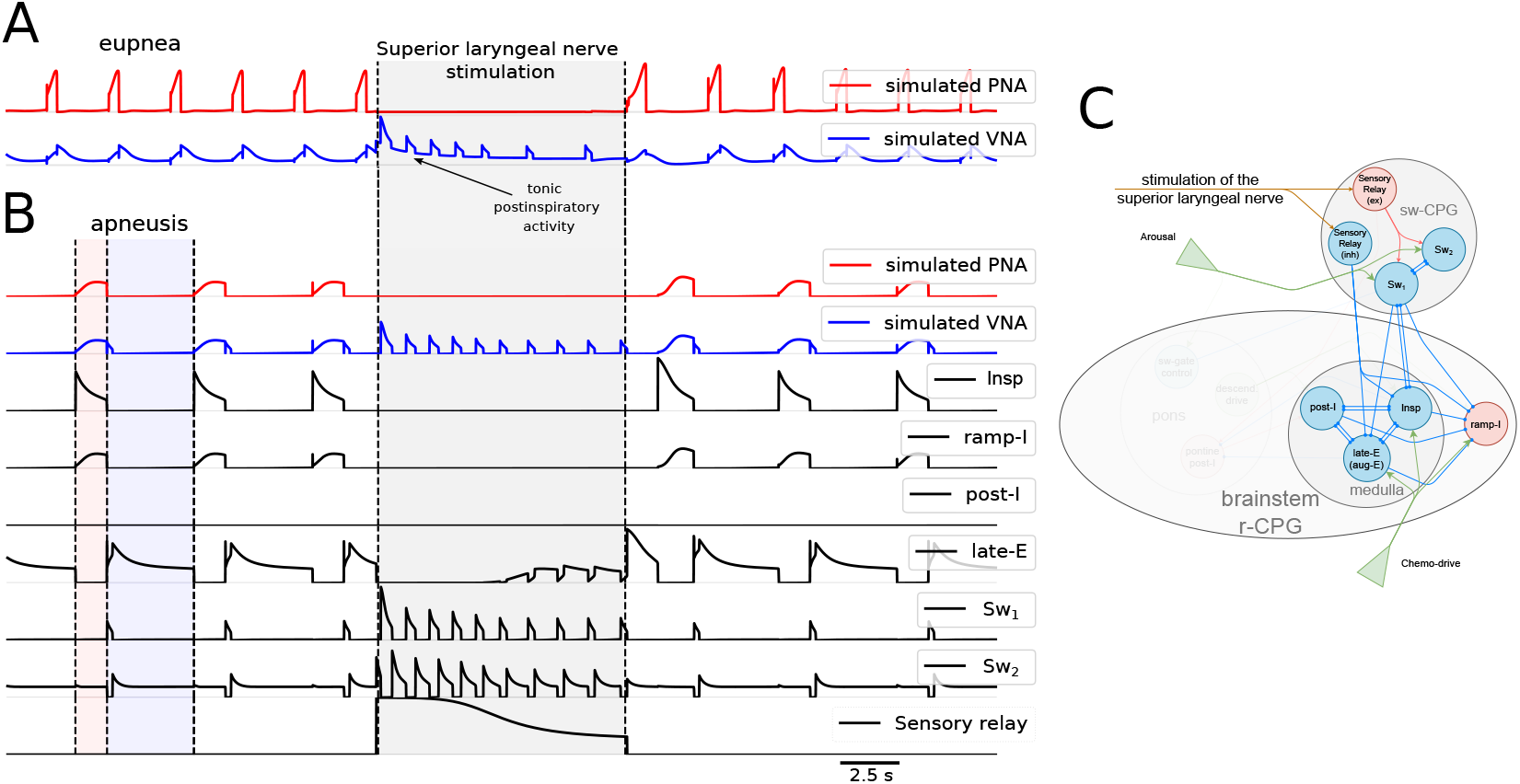
Simulation of the effect of pontine transection on the breathing-swallowing coordination. The lesioned circuit is depicted in panel (C), with all the pontine populations, as well as the pontine drive, completely removed. The simulated phrenic and vagus nerve activity and the average membrane potential traces of neuronal populations are depicted in (B). The network demonstrates an apneustic two-phased regime, lacking postinspiratory activity in its output. The 10 s sensory stimulation triggers the sequential swallowing response, although the concomitant glottal closure is absent. The simulated pontine lesion has also alleviated inhibition of premotor swallowing neurons, and the latter become active throughout the respiration, resembling the spontaneous activity observed in the experiments. To highlight the changes in the respiratory rhythm, panel (A) demonstrates the traces of the simulated phrenic and vagus nerve activities in the intact network. See text for more details.

However, since our model is noiseless and deterministic, the simulated spontaneous swallows are manifested as a regular, respiratory-entrained activity, an effect of which was less obvious in experimental studies of pontine inhibition but can be seen during autoresuscitation of the brainstem networks (Bautista et al. (2014a)).

### 3.3 Model validation: phase response curve

Experimental data obtained in a perfused brainstem preparation of a rat showed that the application of short train (20Hz, 250ms) electrical stimulation of the SLN caused specific changes in the respiratory cycle (Fig. 1, Fig. 8). When the stimulation of the SLN was delivered in the inspiratory phase, ongoing PNA was terminated; with short latency, a single swallow was initiated, and the following expiratory interval was shortened (Horton et al. (2018)). The stimulation during the late expiratory phase also evoked a single swallow, however, the stimulation has delayed the onset of the next inspiratory discharge. The effects of the short stimulation of the SLN have been summarised in the phase response curve (PRC), which became a standard tool for analysis of neural oscillations (Galán et al. (2005), Izhikevich (2007)). We observed a good agreement between the PRC extracted from the model with the experimentally acquired curve (Fig. 8).

**FIGURE 8.**
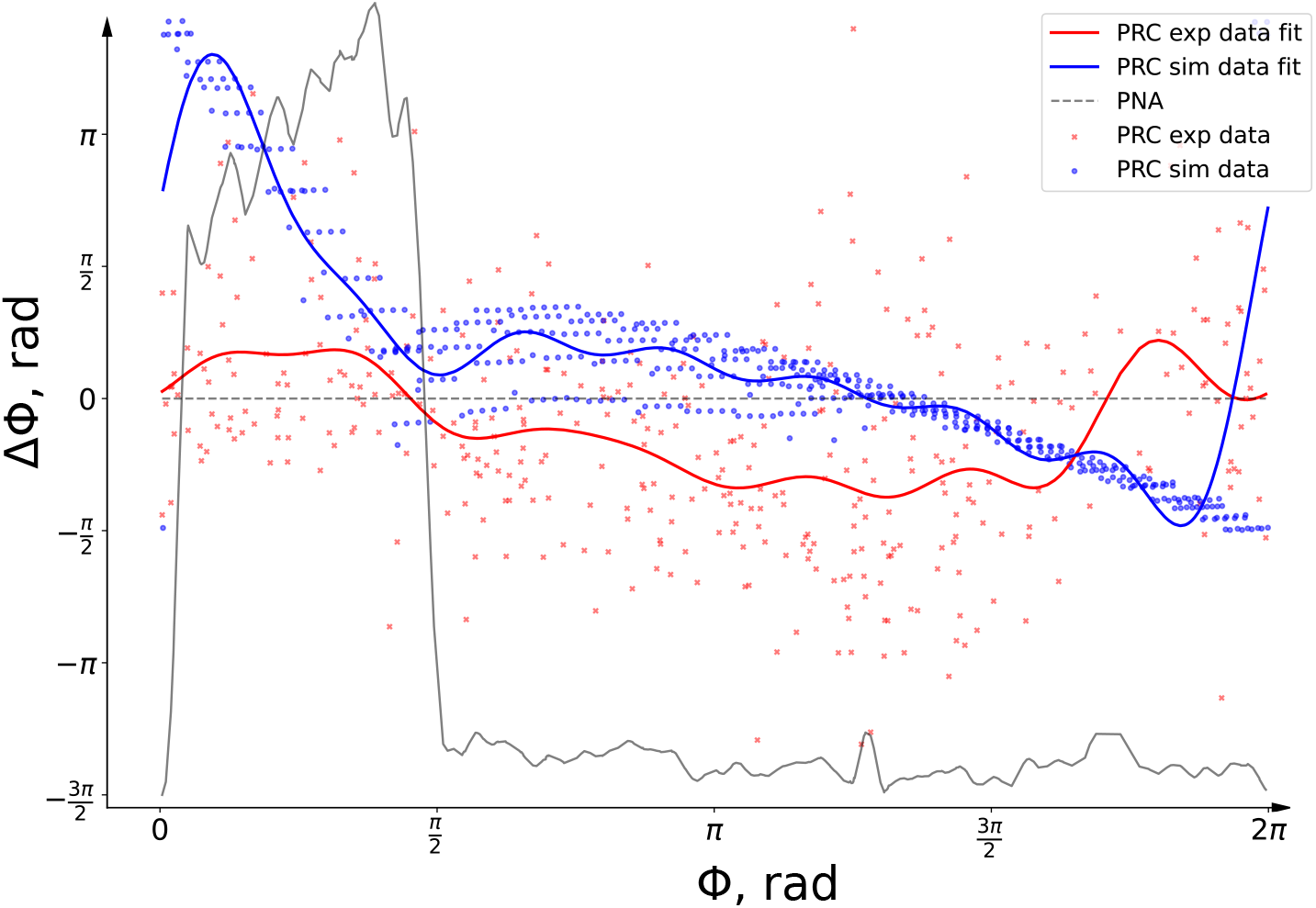
A short stimulation of the superior laryngeal nerve (SLN) elicits a single swallow, which interferes with the respiratory cycle. The figure demonstrates the quantitative comparison of this dynamic in the form of the phase resetting curve (PRC). The PRC estimated from the experimental data (250 ms stimulation, red crosses and red curve) is compared with the PRC calculated from the model simulations (100 ms stimulation, blue circles, blue curve). The PRC calculated from the model qualitatively matches the experimentally obtained curve. The PRCs demonstrate the delay of the next inspiration when the stimulation is applied during the late expiration (the region with the negative phase shift). Both PRC curves capture the effects of a short SLN stimulus application during the inspiration: termination of the inspiration and the new burst of inspiratory activity shortly after (positive phase shift region on the figure).

### 3.4 Model predictions: effects of variation of drives, intrinsic properties and the synaptic connectivity underlying breathing-swallowing interaction

We performed multiple numerical simulations to establish how variations of drive, intrinsic properties, and synaptic connections alter the interaction between the swallowing and the respiratory CPGs. While evaluating the effects of parameter variation, we focused on three aspects (see Fig. 9): the number of swallows during the 10s SLN stimulation (green dots), the occurrence of the spontaneous swallows (blue-hatched regions), and the presence of single swallows following a 100ms SLN stimulation (red-hatched regions).

**FIGURE 9.**
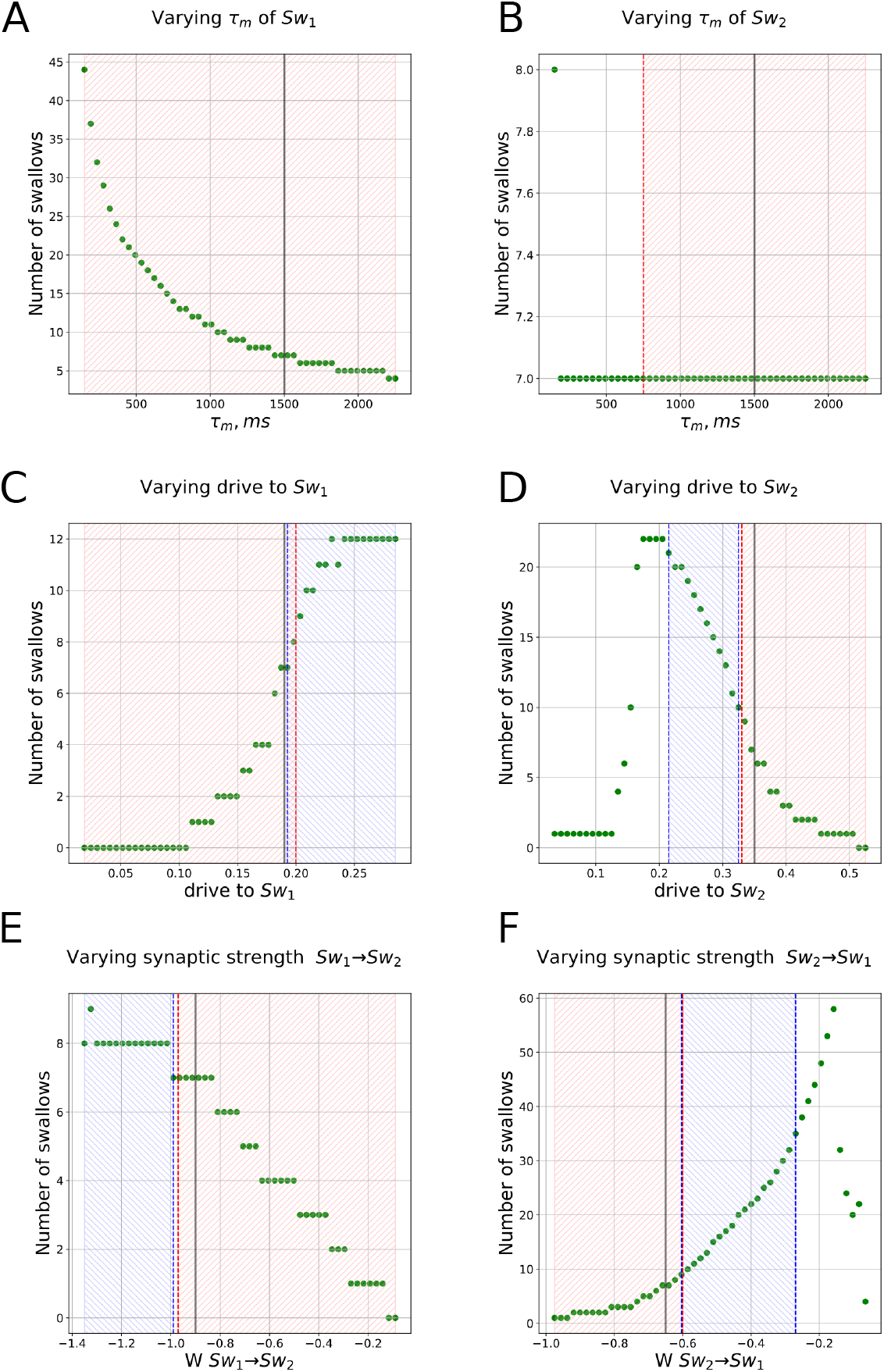
The control of the swallowing: variation of the parameters of the sw-CPG. The number of swallows is indicated by green circles. Blue hatched regions represent the regions where spontaneous swallows occur. The red hatched regions highlight the parameter-regions where the short SLN stimulus evokes a short latency single swallow. The normal value of the parameter is depicted by the grey vertical line. See text for the details.

Varying the spike-frequency adaptation constant of the ‘Sensory Relay’ neurons causes a change in the number of swallows: lower values of time-constant *τ*_*m*_ resulted in the faster spike-frequency adaptation, causing fewer swallows. In our model, the spike-frequency adaptation constant of the ‘Sensory Relay’ neurons is responsible for the gradual decrease of the swallowing frequency at the end of the stimulation.

The variation of spike-frequency adaptation constant *τ*_*m*_ of the ‘Sw_1_’ neurons has a drastic effect on sequential swallowing (see Fig. 9 A). However, for the lower values (*τ*_*m*_ < 500 ms) the unphysiologically high number of swallows (reaching up to 45 swallows during a 10s input simulation) is achieved through unphysiologically short swallow burst durations. The drive to ‘Sw_1_’ neurons controls the presence of both single and spontaneous swallows (Fig. 9 C): for the drives greater 0.19, no single swallows could be triggered, and the network demonstrates constant swallowing activity throughout the respiration. The width of individual swallows in the sequential swallowing during the 10s SLN stimulation as well as the ability to evoke a single swallow is also controlled by the drive to the ‘Sw_2_’ population (Fig. 9 D). In addition, low drives to ‘Sw_2_’ neurons (< 0.21) cause a disinhibition of ‘Sw_1_’ neurons, leading to the generation of spontaneous swallows and interference with the respiratory regime. A larger excitatory drive to ‘Sw_2_’ neurons (> 0.5) abolished the sequential swallows during the 10s SLN stimulus, however, single swallows triggered by 100ms SLN stimulation could still be evoked (Fig. 9 D).

Variation of the inhibition from ‘Sw_2_’ to ‘Sw_1_’ revealed that the number of swallows in the sw-CPG grows when the inhibition has lower absolute values (from −0.6 to −0.2), reaching as high as 60 swallows, however, the width of these swallows was unphysiologically short (Fig. 9 F).

Insufficient activation of ‘Sw_2_’ by the ‘Sensory Relay’ neurons during the SLN stimulation leads to the absence of the single after-stimulus swallows. The absence of the single swallows is also observed in case of increased excitation to the ‘Sw_1_’ neurons coming from the ‘Sensory Relay’ neurons. The lack of the swallowing response after the stimulus strongly correlates with the timing of the swallows during the long stimulation: if the short SLN stimulus immediately produces a swallowing response, and no swallowing response is observed after the stimulus, there is no delay in sequential swallowing during the 10 seconds stimulation.

In our model, the pathological inspiratory breakthroughs occur when the inhibition from the ‘Sensory Relay’ neurons to inspiratory neurons (‘Insp’) is not sufficient (Fig. 10, *W*_*SR*→*Insp*_ = *W*_*SR*→*RampI*_ = −0.22 instead of −0.4). These breakthroughs occur towards the end of the evoked sequential swallowing response. In our simulations, the inspiratory breakthroughs are biphasic since the inspiratory activity is under strong inhibition from the ‘Sw_1_’ neurons, and an inspiratory breakthrough is split into two parts by the generated swallows.

**FIGURE 10.**
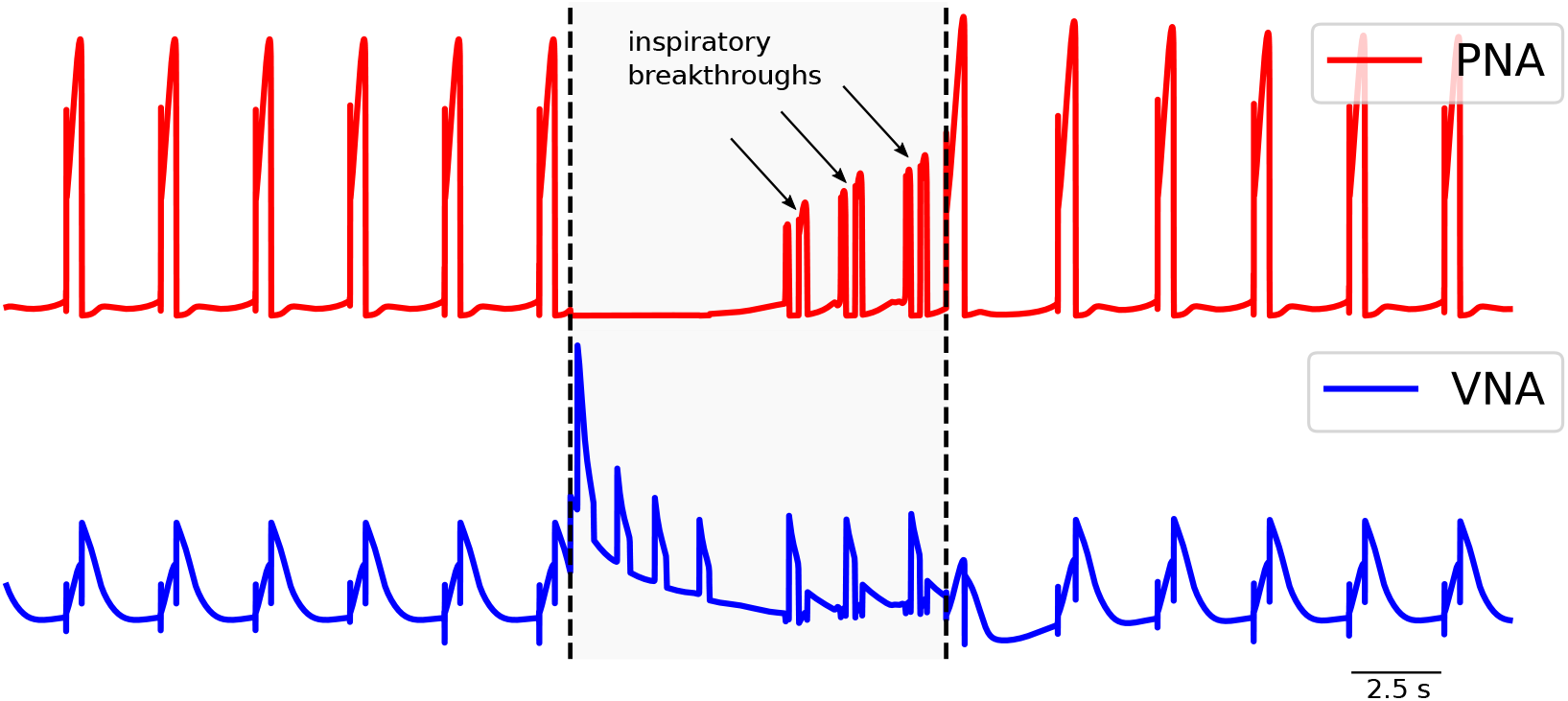
The simulated pathological activity in the phrenic nerve caused by the weakened sensory inhibition of the inspiratory neurons, both ‘Insp’ and ‘ramp-I’. In the simulation, the strength of the sensory inhibition *W*_*SR*→*Insp*_ and *W*_*SR*→*RampI*_ is −0.22 instead of −0.4 in the intact network. The simulated inspiratory breakthroughs are biphasic, since the inspiratory neuronal populations are inhibited by the ‘Sw_1_’ group, and the latter becomes active during the inspiratory breakthrough.

In the model, a delayed and insufficient swallowing response to the simulated 10s SLN stimulation is linked to reduced strength of sensory inputs to sw-CPG (Fig. 11, *W*_*SR*→*RampI*_ = 0.052, whereas the normal value is 0.12). Reduced sensory excitation of ‘Sw_1_’ neuron population results in a greater latency of the first swallow (400 ms vs. 190 ms) and diminishes the number of the evoked sequential swallows, resembling the pathological swallowing response in patients with deglutition dysphagia (Daniels and Foundas (1997)).

**FIGURE 11.**
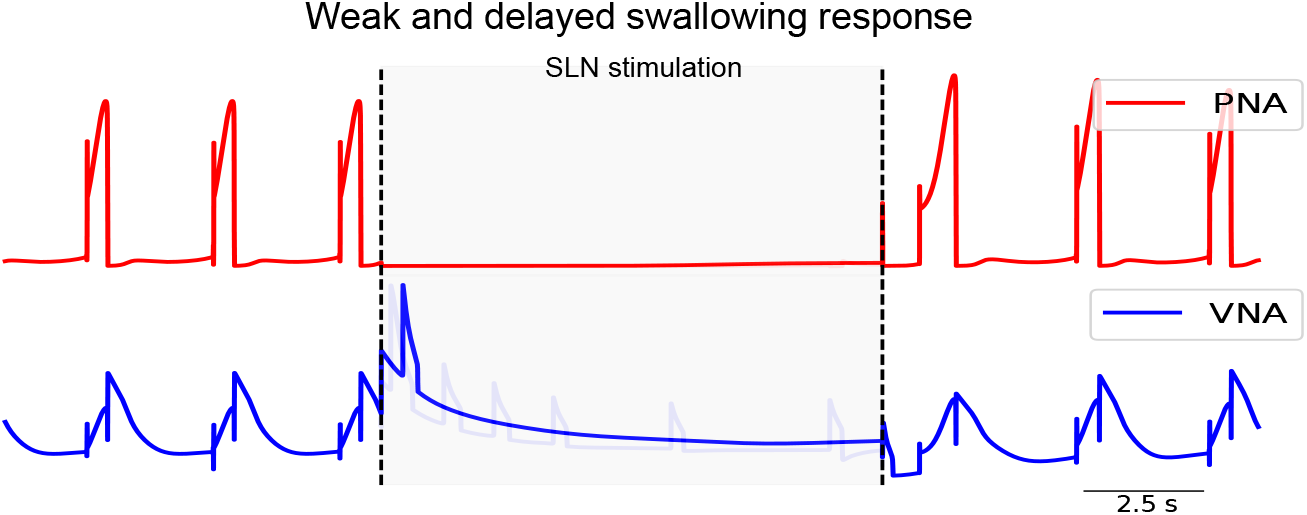
The delayed and weakened swallowing response in the model of the r-CPG-sw-CPG network reflected in the responses of phrenic and vagus nerves to the long SLN stimulation. For the simulation, the parameter *W*_*SR*→*RampI*_ = 0.052, whereas the corresponding value in the intact network is 0.12. The transparent traces of the normal swallowing response are shown in the background for comparison.

## 4 DISCUSSION

Despite multiple research efforts to unravel the connectivity underlying respiratory and swallowing behaviours, a sufficiently detailed model of breathing-swallowing interaction unifying disparate experimental data has not yet been established (Horton et al. (2018)). In the present study, we developed a computational model of the interaction between the breathing and swallowing CPGs that can reproduce three-phase eupneic breathing, distinct swallowing response to the short and long SLN stimulation, and the effects of pontine lesion on the interaction between the two circuits. To describe the dynamics of the neural populations, we utilised simplistic equations underlying the Matsuoka neural oscillator: only the intrinsic property mediating spike-frequency adaptation is incorporated into the equations, allowing to focus on the connectivity. The external tonic drives, mutual inhibition and the spike-frequency adaptation were enough to account for the experimental data listed above. The model generates testable predictions which may further guide the experiments aiming at unravelling the network connectivity.

### 4.1 Network dynamics and the intrinsic properties

The current model of the interaction between the swallowing and breathing CPGs builds on the previous computational models of the respiratory circuit (Molkov et al. (2017)). The preceding models of the respiratory circuit have utilised either conductance-based models, constructed from the Hodgkin-Huxley (HH) neurons (Smith et al. (2007), Molkov et al. (2013), Rybak et al. (2004)) or activity-based models, whose dynamics were described in terms of average membrane potentials of the interacting neuronal populations (Rubin et al. (2009), Molkov et al. (2014)). Although utilising much fewer network units than the conductance-based models, the activity-based models still were able to capture the key dynamical behaviours of the respiratory circuit (Molkov et al. (2017)). However, being inherited from the conductance-based models, the preceding activity-based models utilised detailed HH-style description of the slow ionic currents, which require multiple intrinsic parameters to be extrapolated (Del Negro et al. (2018)). In contrast with the previous efforts, we discarded the description of the biophysical properties in the HH-style in favour of simpler equations underlying the Matsuoka neural oscillator. This conceptual difference resulted in more succinct model description, easier reproducibility and allowed for more straightforward analysis. At the same time, the model is still able to capture the physiologically relevant three-phase rhythmic breathing and effects of pontine transection (Smith et al. (2007)) and distinct swallowing responses to the stimulation of the SLN.

Interestingly, it has been established that different ion channel compositions may lead to the same qualitative dynamical properties of a neuron (Schulz et al. (2006), Marder and Bucher (2007), Goaillard et al. (2009), Golowasch (2014), Marder et al. (2015)). This supports our claim that for understanding the network dynamics, a description of the circuit dynamics on a population level is more important than a description of intrinsic properties associated with ionic currents. However, characterising neuronal ion channel composition is still indispensable for the models aiming to predict the effects of specific drugs.

### 4.2 The functionality of the swallowing CPG

Previous computational efforts of the respiratory CPG have featured the models incorporating both intrinsically bursting neurons and mutually inhibiting populations with the spike-frequency adaptation property (SFA-mediated dynamics). Following the discovery of the inspiratory related intrinsically bursting neurons in the Pre-Bötzinger (Smith et al. (1991)), the ion channel kinetics have been extensively characterised for these neurons (Butera Jr et al. (1999), Ptak et al. (2005)). Previous models have incorporated detailed description of these inspiratory bursting neurons, however, recent computational studies report that the contribution of the bursting neurons in the intact respiratory network may be masked by the synaptic mechanisms (Rubin and Smith (2019), Wang and Rubin (2020), Tolmachev et al. (2018), Guerrier et al. (2015)). In this study, we assume that the bursting properties of the neurons in the PreBötC are not crucial for the interaction between the intact breathing and swallowing circuits. Thereby, the present model features the respiratory CPG without intrinsically bursting neurons, relying solely on the SFA-mediated dynamics.

The same dichotomy of the primary source of the rhythmogenesis extends to the swallowing circuit. Little is known about the dynamics of the swallowing network, although it is likely that both bursting and reciprocal inhibition contributes to the dynamics of the swallowing (Jean (2001), Steuer and Guertin (2019)). Similar to the respiratory network (Rubin and Smith (2019)), relying on multiple dynamical mechanisms allows the swallowing network to function more robustly (Marder et al. (2015)). Nevertheless, for the concise description of the model, we hypothesised that the functionality of the swallowing CPG is based on a half-centre oscillator (HCO) and SFA-mediated dynamics. The swallowing HCO hypothesis is not new and was previously suggested in a computational study by Bolser et al. (2015), however, the authors did not proceed with the analysis of the circuit. The swallowing HCO hypothesis is sufficient to explain a sequential swallowing pattern with a declining frequency of swallows during the 10 s SLN stimulation. Moreover, Bautista and Dutschmann (2014) have shown that the local *GABA*_*A*_ receptor blockade in the NTS causes a marked disturbance of the swallowing response to water stimulation, which corroborates the hypothesis. However, the evidence is still insufficient to state the organisation of the swallowing CPG decisively: further research is needed to determine the precise source of the sequential burst of activity in the swallowing circuit.

Ubiquitous in CPG, and HCOs in particular (Hooper (2001), Marder and Bucher (2001), Bucher et al. (2015), Nagornov et al. (2016)), interneurons often demonstrate intrinsic properties underlying postinhibitory rebound. The presence of the intrinsic properties leading to a postinhibitory rebound dynamic in the swallowing circuit was hypothesised yet in (Jean (2001)), and low threshold-activated calcium channels *I*_*T*_(*Ca*) were reported to mediate this property in the swallowing-related neurons in the nucleus of the solitary tract (Tell and Bradley (1994), Champagnat et al. (1986), Jean and Dallaporta (2006)). We hypothesise that the postinhibitory rebound dynamics in the swallowing CPG may underlie the short-latency single swallowing response triggered by a transient SLN stimulation. The short latency in the response can also arise from slow synaptic NMDA-mediated excitation (Jean (2001), Tell and Jean (1993)), or from early transient outward potassium current *I*_*KA*_ (Jean (2001), Tell and Jean (1991)). In our model, however, a generation of a single short-latency swallow is attributed to dynamics of a whole swallowing HCO (see Sec. 3.1), rather than to the intrinsic properties of specific neurons: the swallowing HCO operates in a quasi-bistable regime with the premotor swallowing neurons demonstrating the transient after-stimulus activity, which we attributed to a single swallow (see Fig. 6 A). Thus, the single swallowing response may be mediated by the dynamics of the swallowing HCO, relying solely on reciprocal inhibition and an SFA property. Although multiple mechanisms may underlie the generation of the single swallowing response, this is the first computational study that addresses this issue.

Regardless of the precise mechanism underlying the single swallowing response, a short stimulation of the SLN during the inspiration reliably evokes a swallow which terminates the inspiratory activity, and the inspiration is resumed with some delay shortly after the swallow has terminated (Lewis et al. (1990)). The SLN stimulation during the late stages of expiration introduces a delay in the onset of the next inspiratory discharge (Jean (2001), Jean and Dallaporta (2013)). This likely occurs because of the mutual inhibition between the inspiratory and the swallowing-related neurons (Horton et al. (2018), Saito et al. (2003), Saito et al. (2002)). We tested how well the mutual inhibition accounts for the experimental observations by comparing the model-derived and experimental phase response curves. The qualitative match of two curves is further evidence for the mutual inhibition between the premotor swallowing and the inspiratory neurons (Del Negro et al. (2018)).

### 4.3 Mediation of the glottal closure reflex and the swallowing breath-hold

Multiple experiments established that the Kölliker-Fuse area (KF) in the dorsolateral pons contributes to the activity of the vagal nerve during normal breathing, modulating the pattern of the expiratory airflow via respiratory-related partial laryngeal adduction (Harding (1984), Dutschmann and Paton (2002b), Dutschmann and Herbert (2006), Browaldh et al. (2016)). The neurons in the KF also contribute to the swallowing-related laryngeal adduction (Dutschmann et al. (2014), Browaldh et al. (2016), Sun et al. (2011)) during the stimulation of the SLN, projecting to the expiratory laryngeal motor neurons in the nucleus ambiguus (Nunez-Abades et al. (1990), Jordan (2001)). It may be hypothesised that the swallowing-related laryngeal adduction may also originate from the medullary postinspiratory neurons in the BötC (Sun et al. (2011)).

However, the inhibition of the BötC neurons does not abolish swallowing-related glottal closure (Sun et al. (2011)), suggesting that the postinspiratory neurons in the medulla are not the primary component in the mediation of the laryngeal adduction reflex attributed to swallowing (Bautista et al. (2014b)). Incorporating these experimental facts into a model, we introduce the pontine neurons (‘pontine post-I’) as the only mediator of swallowing related laryngeal adduction. In our model, removal of this population causes the absence of laryngeal adduction reflex in response to the SLN stimulation, consistent with the experimental data (Bautista and Dutschmann (2014), Browaldh et al. (2016)). Nevertheless, some contribution to the glottal closure reflex may originate from the swallowing circuit (DSG/VSG) itself (Jean (2001), Sugiyama et al. (2011), Sun et al. (2011), Bautista and Dutschmann (2014)).

In the intact network, the long stimulation of the SLN evokes the glottal closure reflex safeguarding the swallows (Jean (2001)), while at the same time the SLN stimulation suppresses the inspiratory activity, causing swallowing breath-hold. The experimental evidence suggests that the swallowing breath-hold is governed separately from the glottal closure after the level of the NTS (Hiss et al. (2003), Bautista et al. (2014b), Sun et al. (2011)). To account for this fact, we hypothesised that, while the laryngeal adductor response is mediated by pontine neurons, the sensory relay neurons directly inhibit the inspiratory neurons in the medulla, hence the two behaviours are carried out by the different pathways. Contrary to our hypothesis, the experimental data indicate that the breath-hold is mediated by the indirect inhibition of the inspiratory neurons: the sensory relay neurons in the NTS provide excitation to the BötC postinspiratory neuronal population, which in turn inhibits the inspiratory neurons in the PreBötC (Sun et al. (2011)). However, implementing an indirect inhibition of the inspiratory neurons in our model resulted in unphysiologically long inspiratory activity after the SLN induced swallowing breath-hold (data not shown).

Our computational model predicts that insufficient swallowing breath-hold response causing the inspiratory breakthroughs results from the deterioration of the underlying connectivity between the NTS and the BötC. In clinical practice, such inspiratory breakthroughs are associated with aspiration, which is discussed in the next section.

### 4.4 Pathological breathing-swallowing interaction and neurologic dysphagia

Swallowing responses can be classified according to the timing at which the response is carried out. The following classes of the swallowing responses can be distinguished: the swallowing initiated in the inspiratory phase (I-Sw-I), the swallowing during the expiration which is followed by the additional expiration phase (E-Sw-E), and the swallowing occurring at the late stage of expiration, directly followed by the inspiration (Sw-I). In clinical practice, I-Sw and Sw-I patterns are associated with increased risk of aspiration (Yagi et al. (2017)), which is an aberrant behaviour of high relevance for the clinical pathology.

Our model allows to hypothesise that the abnormal I-Sw pattern may be associated with decreased inhibition between the premotor swallowing and medullary inspiratory neurons and a concurrent deterioration of the projections from the sensory relay to the respiratory neurons, which normally function to suppress the inspiratory discharge.

The other pathological swallowing pattern, Sw-I, is associated with an inadequate delay of the swallowing response (Yagi et al. (2017)). Our model indicates that the pathological increase in swallowing latency may be caused either by insufficient excitation of the premotor swallowing neurons, incomplete swallowing apnoea, or inappropriate inhibition of premotor swallowing neurons by pontine neurons. The model also allows to speculate that pathological increase of drive to the sensory NTS neurons projecting to the medulla could mediate obstructive sleep apnoea or be involved in sudden infant death syndrome.

Thus, our model can predict network dysfunction underlying aspiration which often causes fatal pneumonia in the elderly (Finiels et al. (2001), Easterling and Robbins (2008)), patients with stroke (Leslie et al. (2002)), chronic obstructive pulmonary disease (Gross et al. (2009), Nagami et al. (2017)), Parkinson’s (Gross et al. (2008)), and Alzheimer’s diseases (Seçil et al. (2016)). The experimental evidence further supports the derived predictions: both patients and transgenic mouse models of neurological diseases demonstrate severe neurodegeneration in NTS and the KF circuitry, for instance, caused by the accumulation of misfolded tau-proteins (Rüb et al. (2001), Dutschmann et al. (2010), Menuet et al. (2011)).

## 4.5 Conclusion

In the present study, we present a first-computational model that addresses the network connectome for the mediation and coordination of swallowing and breathing. The model solidifies the previous experimental data, provides new predictions, and thereby guides further experiments, which are necessary to distinguish between the possible connectivity topologies. Although the model is a first step in the iterative process of understanding the underlying connectivity, it elucidates the possible mechanisms of breathing-swallowing discoordination underlying the aspiration and neurologic dysphagia and lays the foundation for further computational-modelling research on clinically relevant breathing-swallowing pathologies.

## 5 APPENDIX

### 5.1 The modeling parameters

## Abbreviations

CPG: central pattern generator
HCO: half-centre oscillator
SFA: spike frequency adaptation
SLN: superior laryngeal nerve
VN: Vagal Nerve
PN: Phrenic Nerve
NTS: Nucleus of the solitary tract
KF: Kölliker-Fuse

## Notes

### Competing Interest Statement

The authors have declared no competing interest.

